# Context-dependent behavioral plasticity compromises disruptive selection on sperm traits in squid

**DOI:** 10.1101/2020.08.20.258988

**Authors:** Noritaka Hirohashi, Noriyosi Sato, Yoko Iwata, Satoshi Tomano, Md Nur E Alam, Oscar Escolar, Fernando Ángel Fernández-Álvarez, Roger Villanueva, Lígia Haselmann Apostólico, José Eduardo Amoroso Rodriguez Marian

## Abstract

Male animals are not given equal mating opportunities under competitive circumstances. Small males often exhibit alternative mating behaviours and produce spermatozoa of higher quality to compensate for their lower chances of winning physical contests against larger competitors [1]. Because the reproductive benefits of these phenotypes depend on social status/agonistic ranks that can change during growth or aging [2], sperm traits should be developed/switched into fitness optima according to their prospects. However, reproductive success largely relies upon social contexts arising instantaneously from intra- and inter-sexual interactions, which deter males from developing extreme traits and instead favour behavioural plasticity. Nevertheless, the extent to which such plasticity influences developmentally regulated alternative sperm traits remains unexplored. Squids of the family Loliginidae are excellent models to investigate this, because they show sophisticated alternative reproductive tactics (ARTs) by which small males, known as “sneakers”, produce longer spermatozoa and perform extra-pair copulation to attach their sperm packages near the female seminal receptacle (SR). In contrast, large “consort” males have shorter spermatozoa and copulate via pair-bonding to insert their sperm packages near the internal female oviduct [3]. In addition, plasticity in male mating behaviour is common in some species while it is either rare or absent in others. Thus, squid ARTs display a broad spectrum of adaptive traits with a complex repertoire in behaviour, morphology and physiology [3].

---

Here we performed a taxonomically widespread comparison of sperm flagellum length (FL) in squid and cuttlefish species worldwide (Supplementary Table S1). The FL range was 54–156 μm (Fig. 1a) and showed no correlation with adult body size (dorsal mantle length: *t* = −1.106; *P* = 0.28 using a linear model), reproductive activity (gonadosomatic index: *t* = −0.582; *P* = 0.57) or relative testis mass (testicular somatic index, TSI: *t* = −0.310; *P* = 0.76), a widely used indicator for female promiscuity (Supplementary Table S1).

**Figure 1.**
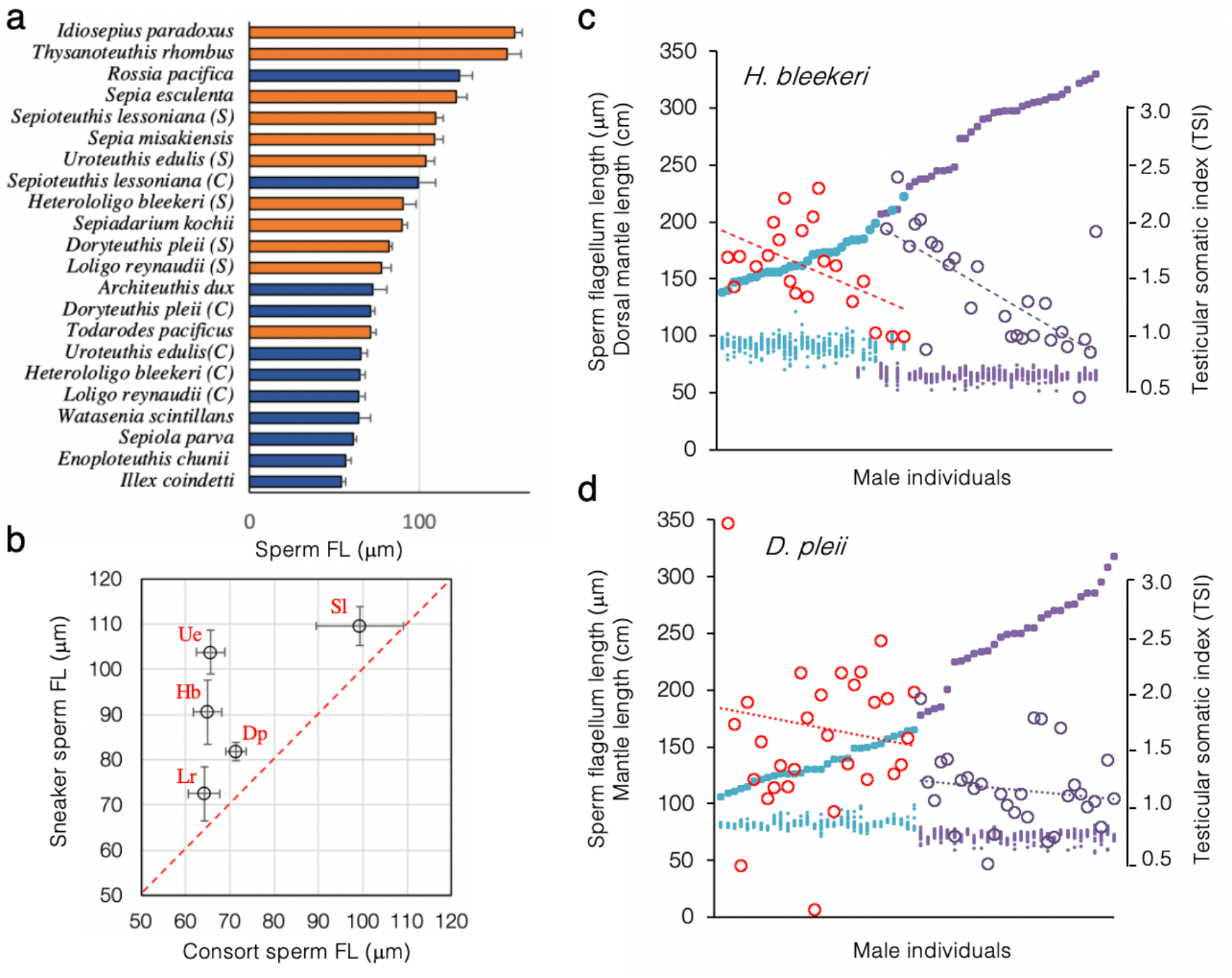
Species comparison of sperm flagellum length (FL) in coleoid cephalopods and detailed comparison of reproductive indices between *H. bleekeri* and *D. pleii.* a, The mean FL arranged by increasing size in 17 coleoid species including 5 species with alternative male types (S, sneaker; C, consort). The colour code indicates the mode of sperm storage, where they are either stored in the female-produced seminal receptacle (*orange*) or in the male-produced sperm package (*blue*). b, Intraspecific sperm dimorphism between sneakers and consorts in five species of loliginid squids. Hb, *H. bleekeri*; Dp, *D. pleii*; Ue, *U. edulis*; Sl, *S. lessoniana*; and Lr, *L. reynaudii.* c–d, Male individual measurements in FL and relative testis mass (testicular–somatic index; TSI) are arranged in order of increasing mantle length. For each male individual in *H. bleekeri* (c) and *D. pleii* (d), ML (*closed circles*), FL values of 20 spermatozoa (*scattered dot*s) and TSI (*open circles*) have been plotted. The colour code represents sneaker males (*cyan* and *red*) or consort males (*purple*).

Female promiscuity and associated sperm competition pressures have been demonstrated as powerful evolutionary drivers for sperm gigantism/ornamentation [4]. However, we found no evidence for sperm competition associated with sperm FL at least within a major clade (Decapodiformes) of cephalopods. For example, the highly promiscuous *Loligo reynaudii* (Supplementary Table S1) has a short flagellum (64.2 ± 3.6 μm), whereas *Thysanoteuthis rhombus*, reported as monogamous [5], has a long flagellum (151.3 ± 8.5 μm; Fig. 1a). In contrast, one parameter that appeared to be correlated with FL was utilization of SR (*t* = −1.106; *P* < 0.01); species (or male type, i.e., sneakers) that use the SR have longer FL (Fig. 1a, orange), whereas those species that do not, have shorter FL (Fig. 1a, blue). This indicates that the SR serves not only as the sperm storage organ but also as the site of mate choice, favouring males with longer spermatozoa. Among *Drosophila* species, FL is associated proportionally with the length of the female genital tract [6], which is widely considered an evolutionary consequence of cryptic female choice (CFC). In cephalopods, SRs are morphologically complex, making it difficult to examine whether CFC or sperm–SR coevolution has occurred. Of note, many species lacking SR have shorter spermatozoa with little variation (Fig. 1a), suggesting that these have the minimal FL required for effective motility. Even if CFC might have driven an evolutionary process for flagellar elongation, any fitness consequence of greater FL remains to be determined in cephalopods.

The trend of close association between FL and SR is also evident in the intraspecific sperm dimorphism of all examined species with ARTs; the FL is longer in sneakers than in consorts (Fig. 1b). However, intraspecific differences in the FL between sneakers and consorts (S/C) varied among species; the difference was substantial in *Uroteuthis edulis* (1.6-fold) and *Heterololigo bleekeri* (1.4-fold), whereas it was small in *Doryteuthis pleii* (1.1-fold) and *Sepioteuthis lessoniana* (1.1-fold). Because these intraspecific, inter-tactic differences would have arisen from disruptive selection by which antagonistic ejaculate traits can evolve, species variation in S/C sperm FL might have arisen from differences in intensity of conflict. We speculate that intensity of tactical conflict can be influenced by the degree of behavioural flexibility. That is, if male mating behaviours are conditional or transformable, ejaculate traits that maximize reproductive fitness of males adopted to each tactic could not be well developed. Based on this assumption, we compared two closely related Loliginidae species, *H. bleekeri* and *D. pleii*, exhibiting contrasting levels in FL dimorphism (Fig. 1b). *H. bleekeri* males exhibit a linear positive relationship in size between spermatophore length and mantle length (ML) and this correlation is discontinuous between sneakers and consorts [7], suggesting that the binary “sneaker-or-consort” fate decision might occur before males become sexually mature. Here, we show further evidence for this hypothesis by revealing ML-based discontinuities in FL and TSI in *H. bleekeri* (Fig. 1c), suggesting that the threshold for dimorphism manifests at earlier stages of development. However, in *D. pleii*, such ML-based discontinuities in sperm FL and TSI are absent (Fig. 1d), in good agreement with previous observations that adult males with intermediate body size (132–178 mm ML) could change their mating tactics flexibly in response to female spawning behaviours [8].Thus, an ontogenetic transition from sneaker to consort might occur within the same individuals [8]. Such flexibility in male mating behaviour was also observed in *S. lessoniana* under experimental conditions where behavioural choice in ARTs is primarily based on the relative body size of the mating partner [9]. These results combined with previous findings by us [7,8] and others [9,10] lead us to speculate that context-dependent behavioural plasticity at mating attenuates disruptive selection on tactic-specific alternative morphs and hence allows the emergence of intermediate phenotypes that could have maladaptive—or moderate—fitness for each mating tactic.

Most squid species grow fast and die within 2 years. Consequently, so-called semelparous reproduction can occur only once and briefly under a high-competition regime. Nevertheless, extremely high degrees of variation in growth rates and body sizes within a population allow individuals to select different strategies for maximizing their mating opportunities, possibly resulting in two different evolutionary trajectories: early ontogenetic decision (*H. bleekeri*) and phenotypic plasticity (*D. pleii*). Interestingly, this difference could be extended to endocrine control, i.e., *H. bleekeri* males can cease their growth after making the decision to become a sneaker, while *D. pleii* can continue to grow even after reaching sexual maturity. Despite the close similarities in life histories and reproductive strategies between these two species, what has driven their diverging evolutionary pathways remains to be determined.

## Supplementary information

### Materials and Methods

#### Sample collection and measurements of individual reproductive and other indices

Locations where animals were collected are: *Illex coindetti* (Vilanova i la Geltrú and Barcelona, Spain), *Sepia misakiensis, Sepiadarium kochii* and *Rossia pacifica* (around Oki Islands, Shimane Pref. Japan), *Sepiola parva* (Kure, Hiroshima Pref. Japan), *Watasenia scintillans, Todarodes pacificus* and *Thysanoteuthis rhombus* (Oki Islands and Sakai-port off, Tottori Pref. Japan), *Sepioteuthis lessoniana* (Tanegashima, Yakushima, Amami Islands, Kagoshima Pref. and Goto Islands, Nagasaki Pref. Japan), *Heterololigo bleekeri* (Tsugaru Strait, Hokkaido Japan)*, Uroteuthis edulis* (Oki Islands and Tsushima Strait, Nagasaki Japan)*, Architeuthis dux* (Wakasa Bay, Kyoto Japan)*, Idiosepius paradoxus* (Chita Peninsula, Nagoya Pref. Japan)*, Enoploteuthis chunii* (Toyama Bay, Ishikawa Pref. Japan), *Sepia esculenta* (Shimane and Tottori Pref. Japan)*, Doryteuthis pleii* (São Sebastião Island, São Paulo Brazil)*, Loligo reynaudii* (Port Elizabeth off, South Africa). The collection methods are either by commercial set-net and trawling fisheries, or luring and net trapping by researchers. Animals obtained from commercial fishing were already dead, therefore dissections and measurements were immediately conducted. All procedures performed in the studies were in accordance with the ethical standards of the Animal Research Committee of Shimane University (ARCSU) and animal experiments were approved by the ARCSU (MA2-2).

#### Sperm size measurement

Sperm flagellum length was measured as described before (Supplementary ref. 5). Briefly, sperm were released from the spermatophores collected from male’s spermatophoric sac or from the spermatangia attached on the female body. Sperm were fixed with 1~4% formaldehyde in seawater and stored at room temperature. Aliquots of formaldehyde-fixed sperm were mounted on a slide-glass which was viewed under a microscope (Nikon TE-2000) with X10-X20 objective lenses. Images were captured with a USB camera and flagellum length was measured with ImageJ software (National Institutes of Health).

#### Statistics

Generalized linear models were used to analyze whether sperm flagellum length was correlated with sperm storage type, GSI, TSI, and ML. All analyses were performed in R 3.4.2 (R Development Core Team 2017).

**Supplemental Table SI.** Surrnmary of reproductive characteristics and other indices of adult individuals in squids and cuttlefishes analyzed in this study.

| species | family | main samping sites | sperm flagellum (μm) |  | n |  | mode of sperm storage | location of sperm storage |
| --- | --- | --- | --- | --- | --- | --- | --- | --- |
|  |  |  | mean | std | male | sperm |  |  |
| *Illex coindetti* | Ommastrephidae | Vilanova i la Geltrú, es | 54.0 | 2.4 | 10 | 200 | spermatangium | near oviduct |
| *Enoplateuthis chunii* | Enoplateuthidae | Toyama Bay, jp | 56.3 | 3.3 | 9 | 135 | spermatangium | near cervical cartilage |
| *Sepiella parva* | Sepiidae | Hiroshima, jp | 60.8 | 1.8 | 5 | 71 | spermatangium* | near oviduct |
| *Watasenia scintillans* | Enoplateuthidae | Shimane/Tottori, jp | 63.9 | 7.4 | 11 | 110 | spermatangium | near cervical cartilage |
| *Loligo reynaudii* (consort) | Loliginidae | Port Elizabeth off, za | 64.2 | 3.6 | 22 | 280 | spermatangium | near oviduct |
| *Heterololigo bleekeri* (consort) | Loliginidae | Tsugaru-Strait, jp | 64.9 | 3.2 | 30 | 600 | spermatangium | near oviduct |
| *Uroteuthis edulis* (consort) | Loliginidae | Oki /Tsushima, jp | 65.7 | 3.3 | 52 | 1030 | spermatangium | near oviduct |
| *Doryteuthis pleii* (consort) | Loliginidae | São Sebastião Island, br | 71.4 | 2.3 | 30 | 600 | spermatangium | near oviduct |
| *Todarodes pacificus* | Ommastrephidae | Oki, jp | 70.9 | 3.3 | 4 | 45 | seminal receptacle | buccal membrane |
| *Architeuthis dux* | Architeuthidae | Wakasa-Bay, jp | 72.5 | 8.4 | 1 | 17 | spermatangium | mantle under skin |
| *Doryteuthis pleii* (sneaker) | Loliginidae | São Sebastião Island, br | 81.8 | 2.0 | 30 | 600 | seminal receptacle | buccal membrane |
| *Loligo reynaudii* (sneaker) | Loliginidae | Port Elizabeth off, za | 77.3 | 5.9 | 14 | 280 | seminal receptacle | buccal membrane |
| *Sepiadarium kochii* | Sepiidae | Oki, jp | 89.8 | 3.3 | 1 | 20 | seminal receptacle | buccal membrane |
| *Heterololigo bleekeri* (sneaker) | Loliginidae | Tsugaru-Strait, jp | 90.5 | 7.2 | 30 | 600 | seminal receptacle | buccal membrane |
| *Sepioteuthis lessoniana* (consort) | Loliginidae | Oki/Tanegashima/Yaku shima/Amami/Goto, jp | 99.3 | 9.8 | 16 | 311 | spermatangium | near oviduct |
| *Uroteuthis edulis* (sneaker) | Loliginidae | Oki /Tsushima, jp | 103.8 | 4.8 | 33 | 750 | seminal receptacle | buccal membrane |
| *Sepia misakiensis* | Sepiidae | Oki, jp | 109.0 | 5.1 | 2 | 29 | seminal receptacle | buccal membrane |
| *Sepioteuthis lessoniana* (sneaker) | Loliginidae | Oki/Tanegashima/Yaku shima/Amami/Goto, jp | 109.6 | 4.3 | 15 | 321 | seminal receptacle | buccal membrane |
| *Sepia esculenta* | Sepiidae | Shimane/Tottori, jp | 121.7 | 5.9 | 5 | 65 | seminal receptacle | buccal membrane |
| *Rossia pacifica* | Sepiidae | Oki, jp | 123.1 | 8.2 | 5 | 50 | spermatangium | near oviduct |
| *Thysanoteuthis rhombus* | Thysanoteuthidae | Oki/Sakai-port, jp | 151.4 | 8.5 | 2 | 55 | seminal receptacle | buccal membrane |
| *Idiosepius paradoxus* | Idiosepiidae | Chita-Peninsula, jp | 156.2 | 4.5 | 12 | 120 | seminal receptacle | buccal membrane |

| species | TSI |  | GSI |  | n | habitat | adult DML range (mm) |  | sexual size dimorphism (difference: mm) | refs |
| --- | --- | --- | --- | --- | --- | --- | --- | --- | --- | --- |
|  | mean | std | mean | std |  |  | male | female |  |  |
| *Illex coindetti* | 2.81 | 0.39 | 6.05 | 0.95 | 20 | neritic-oceanic | 130-140 | 160-170 | female-biased (30) | 1 |
| *Enoplateuthis chunii* | 4.08 | 0.33 | 8.35 | 1.07 | 8 | mesopelagic | 50-100 |  | - | 2 |
| *Sepiella parva* | 3.66 | 0.98 | 6.91 | 1.01 | 13 | sublittoral | <10 |  | - |  |
| *Watasenia scintillans* | 0.10 | 0.05 | 2.36 | 0.61 | 149 | mesopelagic | 45-50 | 50-60 | female-biased (10) | 3 |
| *Loligo reynaudii* (consort) | 1.86 | 0.72 | 3.21 | 0.96 | 159 | neritic | 160-430 | 140-260 | male-biased (170) | 4 |
| *Heterololigo bleekeri* (consort) | 1.50 | 0.47 | 2.64 | 0.68 | 45 | neritic | 160-400 | 120-300 | male-biased (100) | 5 |
| *Uroteuthis edulis* (consort) | 1.93 | 0.47 | 2.80 | 0.48 | 268 | neritic | 155-410 | 160-250 | male-biased (160) |  |
| *Doryteuthis pleii* (consort) | 1.44 | 0.39 | 2.55 | 0.74 | 220 | neritic | 165-320 | 151-175 | male-biased (145) | 6 |
| *Todarodes pacificus* | 3.31 | 0.54 | 6.25 | 1.67 | 16 | neritic-oceanic | 140-230 | 150-300 | female-biased (170) | 7 |
| *Architeuthis dux* | - | - | 0.70 | - | 2 | mesopelagic | 900-1600 | 800-2000 | female-biased (400) | 8, 9 |
| *Doryteuthis pleii* (sneaker) | 1.79 | 0.74 | 2.65 | 0.90 | 67 | neritic | 90-165 | 151-175 | female-biased (10) | 6 |
| *Loligo reynaudii* (sneaker) | 2.68 | 0.79 | 4.28 | 0.92 | 66 | neritic | 130-200 | 140-260 | female-biased (60) | 4 |
| *Sepiadarium kochii* | ND | ND | ND | ND | ND | sublittoral | <20 |  | - | 10 |
| *Heterololigo bleekeri* (sneaker) | 1.32 | 0.30 | 2.88 | 0.56 | 28 | neritic | 140-260 | 120-300 | female-biased (40) | 5 |
| *Sepioteuthis lessoniana* (consort) | 0.53 | 0.24 | 0.88 | 0.31 | 17 | neritic | spring 120-270<br>fall 240-400 | spring 76-375<br>fall 76-325 | spring: female-biased (95)<br>fall: male-biased (75) | 11 |
| *Uroteuthis edulis* (sneaker) | 1.53 | 0.44 | 2.62 | 0.71 | 630 | neritic | 126-175 | 160-250 | female-biased (75) |  |
| *Sepia misakiensis* | 0.90 | 0.16 | 1.80 | 0.50 | 4 | neritic | <65 |  | - | 12 |
| *Sepioteuthis lessoniana* (sneaker) | 0.49 | 0.25 | 0.98 | 0.64 | 26 | neritic | spring 120-270<br>fall 210-360 | spring 76-375<br>fall 76-325 | spring: female-biased (105)<br>fall: male-biased (35) | 11 |
| *Sepia esculenta* | 1.58 | 0.44 | 2.69 | 0.68 | 3 | neritic | 98-196 | 92-192 | male-biased (4) | 13 |
| *Rossia pacifica* | 0.63 | 0.61 | 3.92 | 0.67 | 5 | sublittoral | 45-50 | 68 | female-biased (18) |  |
| *Thysanoteuthis rhombus* | 0.38 | - | 0.71 | - | 1 | mesopelagic | 55-80 | 64-85 | female-biased (5) | 14 |
| *Idiosepius paradoxus* | 6.05 | 1.26 | 9.13 | 1.24 | 102 | neritic | summer 4.2-8<br>winter 7.7-14 | summer 6.9-14<br>winter 11.1-20 | summer: female-biased (6)<br>winter: female-biased (6) | 15, 16 |

es, Spain; jp, Japan; za, South Africa; br, Brazil; \* bursa copulatrix; n, number; TSI, testicular-somatic index GSI, gonadosomatic index DML, dorsal mantle length; refs, references; -, data not available; blue boxes, data were obtained from the references indicated. consort, consort males; sneaker, sneaker males

## Acknowledgements

We thank Mr. Tetsushi Haraguchi (Miyazu Energy Laboratory Aquarium, Japan) for giant squid specimens and Drs. Chuan-Chin Chiao, Chen-Yen Lin and Chih-Shin Chen for the data of *S. lessoniana*. This study was supported by the faculty of Life and Environmental Sciences in Shimane Univ. FÁF-Á. was supported by an Irish Research Council award (Ref. GOIPD/2019/460). JEARM and LHA acknowledge funding provided by AUCANI (grant 967/2018), FAPESP (grant 2017/16182-1), CAPES (Financial Code 001) and CNPq (grants 477233/2013-9 and 142170/2017-8).

## Authors’ contributions

NH conceived this study. All authors collected specimens. NH, NS, YI, MNEA, LHA and JEARM performed experiments and analyzed data. NH wrote the first draft and all other members edited with critical reading.

